# Does microbiome-associated disease affect the inter-subject heterogeneity of human microbiome?

**DOI:** 10.1101/730747

**Authors:** Zhanshan (Sam) Ma, Lianwei Li

## Abstract

*Space* is a critical and also challenging frontier in human microbiome research. It has been argued that lack of consideration of scales beyond individual and ignoring of microbe dispersal are two crucial roadblocks in preventing deep understanding of the heterogeneity of human microbiome. Assessing and interpreting the spatial distribution (dispersal) of microbes explicitly are particularly challenging, but implicit approaches such as Taylor’s power law (TPL) can still be effective and offer significant insights into the heterogeneity in abundance and distribution of human microbiomes. Here, we investigate the relationship between human microbiome-associated diseases (MADs) and the inter-subject microbiome heterogeneity, or heterogeneity-disease relationship (HDR), by harnessing the power of TPL extensions and by analyzing a big dataset of 25 MAD studies covering all five major microbiome habitats and majority of the high-profile MADs including obesity and diabetes. Our HDR analysis revealed that in approximately 10%-17% of the cases, disease effects were significant—the healthy and diseased cohorts exhibited statistically significant differences. In majority of the MAD cases, the microbiome was sufficiently resilient to endure the disturbances of MADs. Furthermore, comparative analysis with traditional DDR (diversity-disease relationship) results is presented. We postulate that HDR reveals evolutionary characteristics because it utilizes the TPL parameter that implicitly characterizes spatial behavior (dispersion), which is primarily shaped by microbe-host co-evolution and is more robust against disturbances including diseases, while diversity in DDR analysis is primarily an ecological-scale characteristic and is less robust against diseases. Nevertheless, both HDR and DDR cross-verified remarkable resilience of human microbiomes against MADs.

**Importance:** It has been argued that lack of consideration of scales beyond individual and ignoring of microbe dispersal are two crucial roadblocks in preventing deep understanding of the heterogeneity of human microbiome. Assessing and interpreting the spatial distribution (dispersal) of microbes explicitly are particularly challenging, but implicit approaches such as Taylor’s power law (TPL) can still be effective. Here, we investigate the relationship between human microbiome-associated diseases (MADs) and the inter-subject microbiome heterogeneity, or heterogeneity-disease relationship (HDR), by harnessing the power of TPL extensions and by analyzing a big dataset of 25 MAD studies covering all five major microbiome habitats and majority of the high-profile MADs including obesity and diabetes. We postulate that HDR reveals evolutionary characteristics because it utilizes the TPL parameter that implicitly characterizes spatial behavior (dispersion), which is primarily shaped by microbe-host co-evolution and is more robust against disturbances including diseases than the traditional diversity-disease relationship (DDR).

## Introduction

The term “heterogeneity” literally means the quality or state of being diverse in character or content. From its literal meaning, heterogeneity may even be considered as the other side of evenness coin or as a proxy of biodiversity. Nevertheless, much of the formal and quantitative studies of heterogeneity in ecology has been performed separately from diversity research. For example, in population ecology, traditionally the terms heterogeneity, aggregation, dispersion, patchiness can be used interchangeably, and all of which were often used to characterize the spatial distribution of biological population. For this reason, the terms *spatial heterogeneity* and *spatial distribution* are often used interchangeably in population ecology; in the former, the spatial information is referred implicitly and, in the later, the spatial information is referred explicitly. Therefore, to avoid confusion, the terms *spatial* heterogeneity (aggregation, dispersion, patchiness, contagiousness) should be more accurate in ecology. Note that *temporal heterogeneity* can also be defined and measured. In the temporal context, temporal heterogeneity is essentially a proxy of (temporal) stability of population or community (Ma 2015, Oh et al. 2016, Li & Ma 2019). In this study, our focus is spatial heterogeneity.

When the concept of heterogeneity is constrained by the spatial arrangement (distribution) or temporal (variation), its difference with diversity or evenness becomes clear. Obviously, traditional diversity concept and its various metrics do not usually deal with spatial arrangement (*e.g*., Chao *et al*. 2014). We are only aware of two exceptions for the lack of spatial information. One is the concept of beta-diversity, which implicitly consider spatial information but is obviously far less convenient in processing spatial information than the heterogeneity concept. Another exception is to do with the concept of dominance (Ma & Ellison 2018, 2019), which is briefly discussed below.

*Spatial heterogeneity* is a fundamental property of any natural ecosystems including human microbiomes. The *spatial distribution* of microbes on or in our bodies certainly possesses far-reaching implications to our health and diseases. *Space* is not only the last, but also a critical frontier in human microbiome research. In ecology, two most important scales that the spatial heterogeneity exhibits are the *population* and *community*. Of course, to display spatial heterogeneity, there must be at least two individuals in the ecosystem, which implies that heterogeneity is a group (cohort, population, community, *etc*) characteristic. Obviously, the concept of heterogeneity is also widely used in other fields of biology (*e.g*., genetic heterogeneity and landscape heterogeneity).

Theoretically, the abundance and distribution of species are among the most fundamental themes of both theoretical population and community ecology. The concept of *heterogeneity,* especially the spatial heterogeneity, is one of the most successful characterizations of species distribution and abundance (Cohen & Meng 2015, Cohen & Saitoh 2016, Ma 2015, Reuman *et al*. 2014, 2017; Taylor 1984, 2019, Taylor *et al*., 1983, 1988). Practically, being able to predict the heterogeneity scaling (change) over space and/or time, and to understand how the scaling is influenced by environmental disturbances such as diseases is of critical significance.

At population scale, aggregation or dispersion are preferred terms and can be used interchangeably with heterogeneity. In population ecology, *aggregation* can be measured quantitatively with Taylor’s (1961) power law, which achieved a rare status of ecological law (Taylor 2019) as further explained below shortly. In community level or multi-species assembly context, we use the term to refer to the *uneven* or *heterogeneous* nature of species abundances among different species within a community and/or between communities. At community scale, community spatial heterogeneity can be quantitatively measured with Taylor’s power law extensions (TPLEs), which were proposed by Ma (2015) *via* extending Taylor’s (1961) power law from population to community level. On the surface, it appears that the community heterogeneity can be considered as the other side of diversity (evenness) “coin.” However, as explained previously, community spatial heterogeneity deals with spatial information, therefore it is necessary to reiterate that the two sides of the same coin may indeed possess different patterns. In other words, the heterogeneity analysis cannot be replaced by diversity analysis, and the former can offer unique insights, which traditional diversity analysis may not.

In microbial ecology, the spatial distribution of microbes (whether it is the distribution of pathogens or symbionts within our body) is, by no means, less important than in the ecology of plants and animals in nature. The spatial distribution of microbes within our bodies or the intersubject heterogeneity (differences) can influence the competition, coexistence, dispersal, and spread of microbes within or between our bodies, and should have far reaching significance to our healthy and diseases. In fact, the term “heterogeneity” has frequently occurred in the mission statements of both the HMP (Human Microbiome Project) and HMP2 or iHMP (integrative HMP) of US-NIH, referring to various aspects of the human microbiome from inter-subject heterogeneity in microbiome composition to the heterogeneity in disease treatment response (HMP Consortium 2012, iHMP Consortium 2019), which highlighted the significance of the heterogeneity in human microbiome research. Nevertheless, methodological studies on the heterogeneity have been relatively rare and studies that explicitly measure the relationship between microbiome heterogeneity and diseases effects are still few. In the present study, we fill this gap by reanalyzing a big dataset of 16s-rRNA metagenomic datasets of 25 MAD (microbiome associated diseases) studies collected from existing literature, which cover all five major human microbiome habitats (gut, oral, skin, lung and vaginal) and include the majority of the high-profile MADs such as IBD, obesity, diabetes, BV, and neuron-degenerative diseases, and therefore of wide representative of the field.

Indeed, the heterogeneity of human microbiome is remarkable. For instance, it was found that no gut bacterial strains of moderate abundance (>0.5% of the microbial community) were shared among all individuals of human twins (Turnbaugh & Gordon 2009, Miller et al. 2019). However, characterizing, quantifying and interpreting the heterogeneity of human microbiomes are rather challenging (HMP Consortium 2012, iHMP Consortium 2019). For another example, Falony *et al*. (2016) collected 503 host factors possibly influencing human gut microbiome across nearly 4000 individuals, but interpreted merely 16.4% of the variation among individuals. Miller et al. (2019) suggested that two roadblocks are responsible for the lack of explicability to the heterogeneity of human microbiome. First, the analysis scale was usually limited to individual subjects (hosts). Second, the microbial dispersal across individual hosts was often ignored. Directly supporting Miller et al. (2019) arguments are rare but certainly exist. For example, Rothschild *et al*. (2018) revealed that whether subjects lived together or not was the strongest predictor of microbiome variance, indicating that transmission (among hosts and between hosts and their environment) could be a critical driver of microbiome heterogeneity. Taylor’s power law, which achieved a rare status of classic laws in macrobial ecology (Taylor 2019) and was recently extended to community ecology of microbes (Ma 2015, Li & Ma 2019, Oh 2016), offers a powerful tool to remove the roadblocks for better understanding the heterogeneity of human microbiomes because of its inherent connections with spatial distribution and dispersion (Taylor 1983, 1984, 2019).

The objective of this study is therefore two-fold. First, we investigate the relationship between the inter-subject heterogeneity and major human MADs, determining whether or not diseases have significant influences on the inter-subject heterogeneity by harnessing the power of TPLEs in measuring community spatial heterogeneity, as explained above. Second, we compare the relationship between the heterogeneity-disease relationship (HDR) explored in this study with an early meta-analysis by Ma *et al*. (2019) on the diversity-disease relationship (DDR), and further explore the relationship between the two relationships (HDR & DDR) and their ecological interpretations. Similar studies on the spatial distribution (heterogeneity) of host, pathogen and their implications to the disease of plants and animals have been investigating routinely for decades, and their significance to plant pathology and husbandry diseases have been well recognized (Noe & Campbell 1985, Real 1996, Emry *et al*. 2011). Therefore, the study we conduct in this article should have rather important significance to the investigation, diagnosis and treatment of the human microbiome associated diseases, similar to the experience in the studies on the diseases of plants and animals.

## Materials and Methods

### The 16s-rRNA datasets of the human microbiome associated diseases

A brief description on the 16s-rRNA datasets from the 25 human MAD (microbiome associated disease) studies is provided in Table S1 of the online supplementary information (OSI). These datasets covered five major human microbiome habitats (gut, oral, skin, lung, and vaginal) as well as milk and semen fluids. They also included the majority of the most common human MADs such as IBD, obesity, diabetes, periodontitis, cystic fibrosis, mastitis, bacterial vaginosis, Parkinson’s disease, schizophrenia. Therefore, the datasets are rather representative to the field of MADs.

### Conceptual comparisons between population and community heterogeneities

Methodologically, the study of heterogeneity is dependent on statistical *variance* and its relationship with statistical *mean* of species abundance, or the so-termed Taylor’s power law (TPL) (Taylor 1961) and its extensions (Ma 2015, Oh 2016) as explained below. The *variance-mean* power law relationship can reveal certain important community characteristics and insights, which the entropy-based diversity measures fail to offer (Ma 2015, Ma & Ellison 2018, 2019, Li & Ma 2019).

To explain why TPL is one of the most effective tools to investigate the population spatial distribution (heterogeneity), it is necessary to recognize that there are three fundamental spatial distribution patterns of any biological populations from microbes to humans; those are: uniform, random and aggregated. The uniform distribution (sensu biologically) means that individuals of population in space are distributed with equal space and the distribution (pattern) can be described with the *uniform distribution* (sensu statistically). The random distribution means that individuals of population in space are distributed randomly (not necessary with equal space distance) and can be described with Poisson statistical distribution. The aggregated distribution (also known as patchy, clumped, contagious, distribution) means that individuals in space are distributed far from random and often in clusters of different sizes. The statistical distribution for the aggregated or heterogeneous spatial distribution are usually rather special, and often follow highly skewed long-tail distributions such as negative binomial, Ades distribution, power law distribution, or log-normal distribution (Perry & Taylor 1985). Indeed, fitting statistical distributions to population abundance frequency is one of the three major approaches for investigating the spatial distribution (aggregation, heterogeneity, or pattern) of biological populations. The second approach to investigate the spatial heterogeneity is the so-termed aggregation (heterogeneity) index approach. In plant ecology, the spatial information is explicit in the form of neighbor distances between plant individuals in a plot (Greg-Smith 1964). In animal and insect ecology, neighbor-distance information is not often convenient to process, but the metrics and models for measuring aggregation/dispersion/heterogeneity still implicitly deal with spatial information. Various aggregation (heterogeneity) indexes were proposed to measure the aggregation (heterogeneity) levels of the spatial distribution of biological populations. The third approach is the so-termed regression modeling, including two regression models. One is Iwao’s (1968, 1977) *m*^*^-*m* linear model (*m*^*^ is the population mean crowding, itself is a measure of aggregation, and *m* is the population mean). Another is Taylor’s (1961) *V-m* power-law (TPL) model (Vand *m* are the variance and mean of population abundance respectively). The advantages of the third type approaches include the following: (*i*) The regression models (TPL & Iwao models) are essentially the synthesis of the heterogeneity index (*V/m* or *m*^*^/*m*) across even larger scales (multiple populations), and therefore can be utilized to characterize the spatial heterogeneity (distribution) of many populations of a species. In other words, it synthesizes the heterogeneities of multiple populations of a species, while the second approach (aggregation) can only characterize the heterogeneity of a single population. (*ii*) The first approach—distribution fitting—can only describe the heterogeneity of a single population, just as the second approach. Although it contains more detailed information on the spatial heterogeneity than the second approach, it also suffers from the same disadvantage as the aggregation index approach.

At the community level, community spatial heterogeneity (CSH) can be similarly defined with the population spatial heterogeneity (aggregation) (PSH) and investigated with similar approaches. Fitting the species abundance distribution (SAD) such as lognormal and log-series distributions can be considered as the counterpart of the first approach (distribution-fitting to population abundances) in population ecology. The community dominance defined by Ma & Ellison (2018, 2019), which extended Lloyd (1967) mean crowding concept, can be considered as the counterpart of the second approach (aggregation index approach) in population ecology. The four TPL extensions (TPLEs) offer the third approach for investigating the community spatial heterogeneity (Ma 2015, Oh 2016), the counterpart of the third approach (TPL) in population ecology. In the present study, we apply the TPLEs for investigating the spatial heterogeneity of the human microbiomes as well as the influences of the microbiome-associated diseases on the heterogeneity.

### Taylor’s power law extensions (TPLEs)

Taylor (1961, 1984) revealed that the population mean (m) and variance (V) of a biological species in space follows power function,

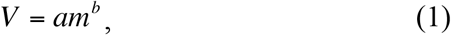

where *m* is the mean population abundance of the species at a specific spatial site, *V* is the corresponding variance, *b* is a species-specific parameter that measures the population aggregation (heterogeneity) degree, and *a* is a parameter primarily influenced by sampling scheme and is of relatively little biological significance. Eqn. (1), known as Taylor’s power law (TPL) in literature, has been verified by numerous field investigations and theoretical analyses and it is considered one of few ecological laws in population ecology (Taylor 2019).

Ma (2015) extended the original TPL from single species population level to community level and introduced four power law extensions (TPLEs). Type-I TPLE was defined to quantify the *community spatial heterogeneity,* and it has the same mathematical form with the original TPL, but with different ecological interpretations, *i.e*.,

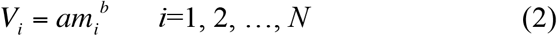

where *m_i_* is the mean species abundance of all species in the *i-th* community (individual host subject) sampled, *i.e*., the mean population size (abundance) per species in the community, *Vi* is the corresponding variance, *N* is the number of total communities (individual subjects) sampled, *a* is similarly interpreted as in the original TPL.

The parameter *b* of Type-I TPLE is a measure for the degree (level) of the community spatial heterogeneity (CSH). The CSH measures the dispersion (difference) in species abundances within a community as well as the scaling of the dispersion (difference) across space (i.e., across communities or sampled individual subjects). The larger the *b*-value is, the higher the community heterogeneity is (Ma 2015, Li & Ma 2019). In the present study, we test whether or not the heterogeneity scaling parameter (*b*) can be significantly influenced by MADs.

Type-III TPLE was defined to assess the spatial heterogeneity of the mixed-species population, with the following formula, which is exactly the same with the original TPL and Type-I TPLE, just with different ecological interpretations:

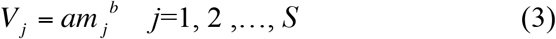

where *mj* is the mean population abundance of a specific species across all communities (individual subjects) sampled, *V_m_* is the corresponding variance, *S* is the number of species in the communities, *a* is sampling scheme related, and *b* measures the spatial heterogeneity of mixed-species.

The concept of mixed species was first discussed by Taylor *et al*. (1983), and it synthesizes species-level spatial heterogeneity across all species. Type-III TPLE for mixed-species spatial heterogeneity synthesizes the dispersion (difference) in population abundance of a single species across space as well as the scaling of the dispersion (difference) across all species. The resultant parameter *b* measures the heterogeneity (aggregation) characteristic of a virtually mixed species, assuming that species identity is irrelevant and can be ignored (Ma 2015, Li & Ma 2019).

We use the log-linear transformations of TPLEs (2-3) to fit them statistically, *i.e*.,

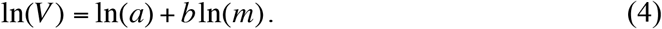

As a side note, Type-II and Type-IV TPLEs were defined to quantify the temporal stability (heterogeneity), but they are not implicated in this study.

Ma (2015) proposed the concept of community heterogeneity critical abundance (CHCA), which was defined with the following formula:

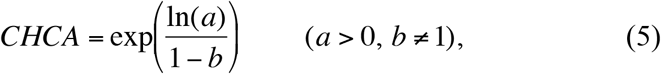

where *a* and *b* are the parameters of TPLE [eqns. (2–3)]. CHCA is a threshold or transition point between the heterogeneous community and regular (completely even) community. At CHCA, the distribution of community heterogeneity should be random. The CHCA is an extension of Ma (1991) population aggregation critical density (PACD) to community scale.

### Randomization tests for the influence of MADs on the spatial heterogeneity

The randomization (permutation) test is performed to determine whether or not the human microbiome associated disease (MADs) possesses statistically significant influences on the scaling (changes) of the spatial heterogeneity. The general procedure for randomization test is referred to Collingridge (2013), and we adapted the general procedure for our testing the TPLE parameters as following five steps. The number of re-samplings for randomization test is set to 1000 times to calculate the *p*-value for determining the significance of differences. The randomization test algorithm used to determine the differences in TPLE scaling parameters is exactly the same as that used in Li & Ma (2019).

## Results

We fitted Type-I & Type-III TPLE models to the healthy (*H*) and diseased (*D*) treatments for each of the 25 MAD studies, respectively. Detailed fitting results were listed in Table S2 (for Type-I TPLE or TPLE-I model) and Table S3 (for Type-III TPLE or TPLE-III model). Fig 1 and Fig 2 illustrated the fittings of Type-I and Type-III TPLE models, respectively. The green lines and points represent for the *H* treatments and the red (and blue) for the *D* treatments.

**Fig 1.**
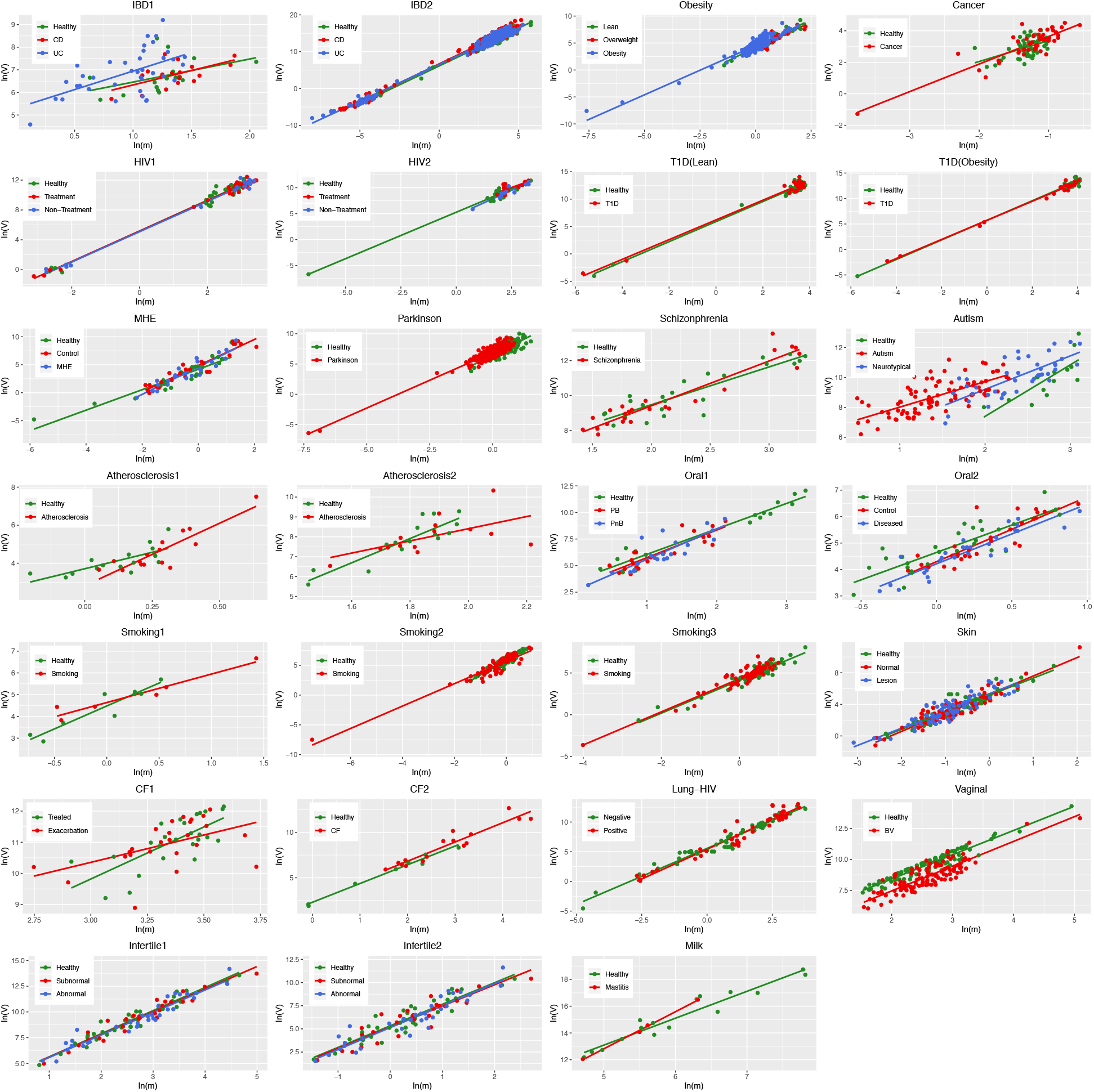
Plots from fitting Type-I Taylor’s power law extension (TPLE-I) for each of the 25 human microbiome-associated diseases (MADs): the heterogeneity parameter (*b*) for measuring community spatial heterogeneity

**Fig 2.**
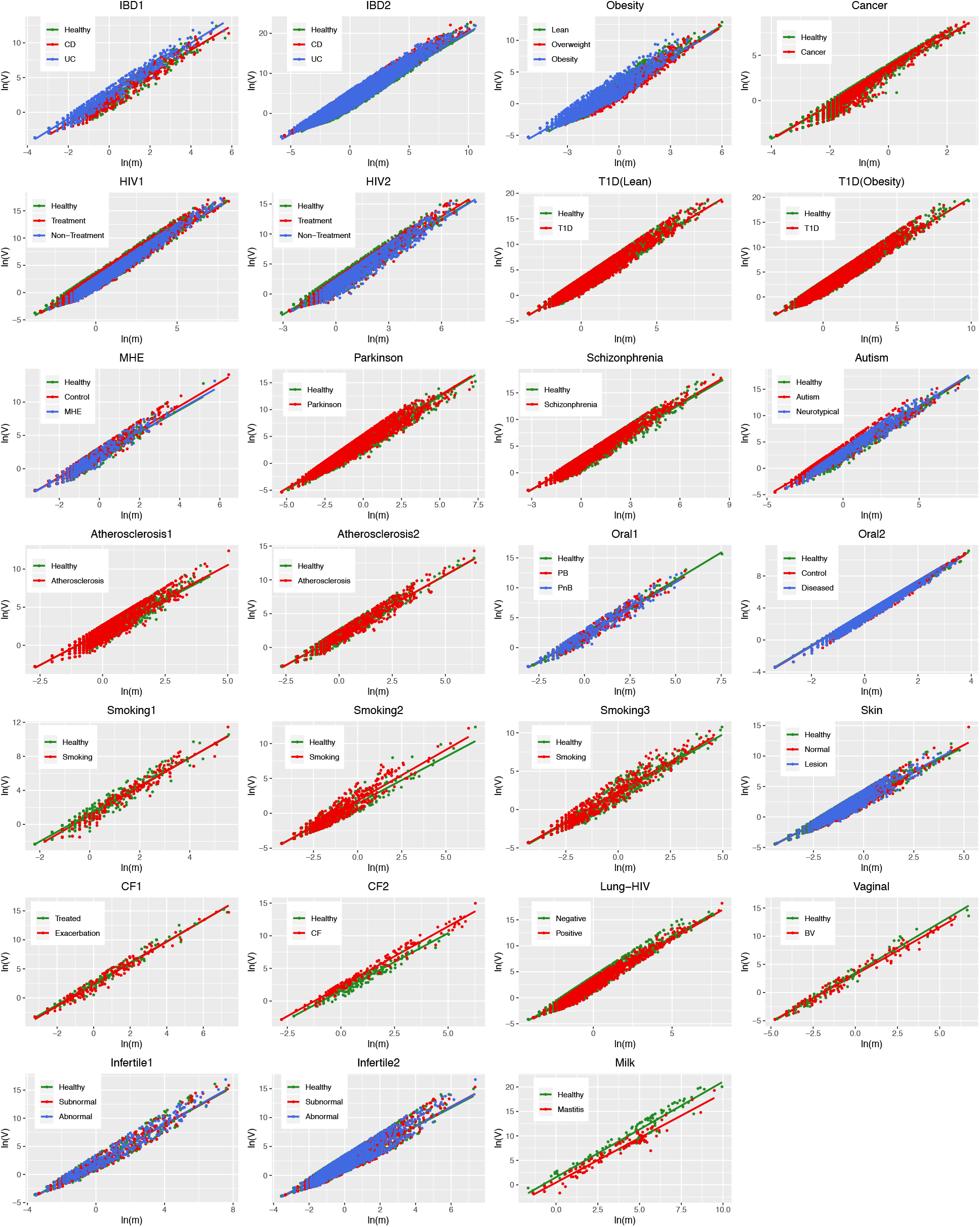
Plots from fitting Type-III Taylor’s power law extension (TPLE-I) for each of the 25 human microbiome-associated diseases (MADs): the heterogeneity (aggregation) parameter (*b*) for measuring mixed-species population heterogeneity (aggregation)

We further performed randomization (permutation) tests to detect the differences in the community spatial heterogeneity (CSH) parameter (*b*) and ln(*a*) between the *H* and *D* treatments (Table S4). The community spatial heterogeneity parameter (*b*) (of TPLE-I) and the mixed-species population aggregation parameter (*b*) (of TPLE-III) were plotted in Fig 3. Fig 3 also marked the cases with significant differences in *b* between the *H* & *D* treatments, which were also displayed in Table S4 (marked with grey shading).

**Fig 3.**
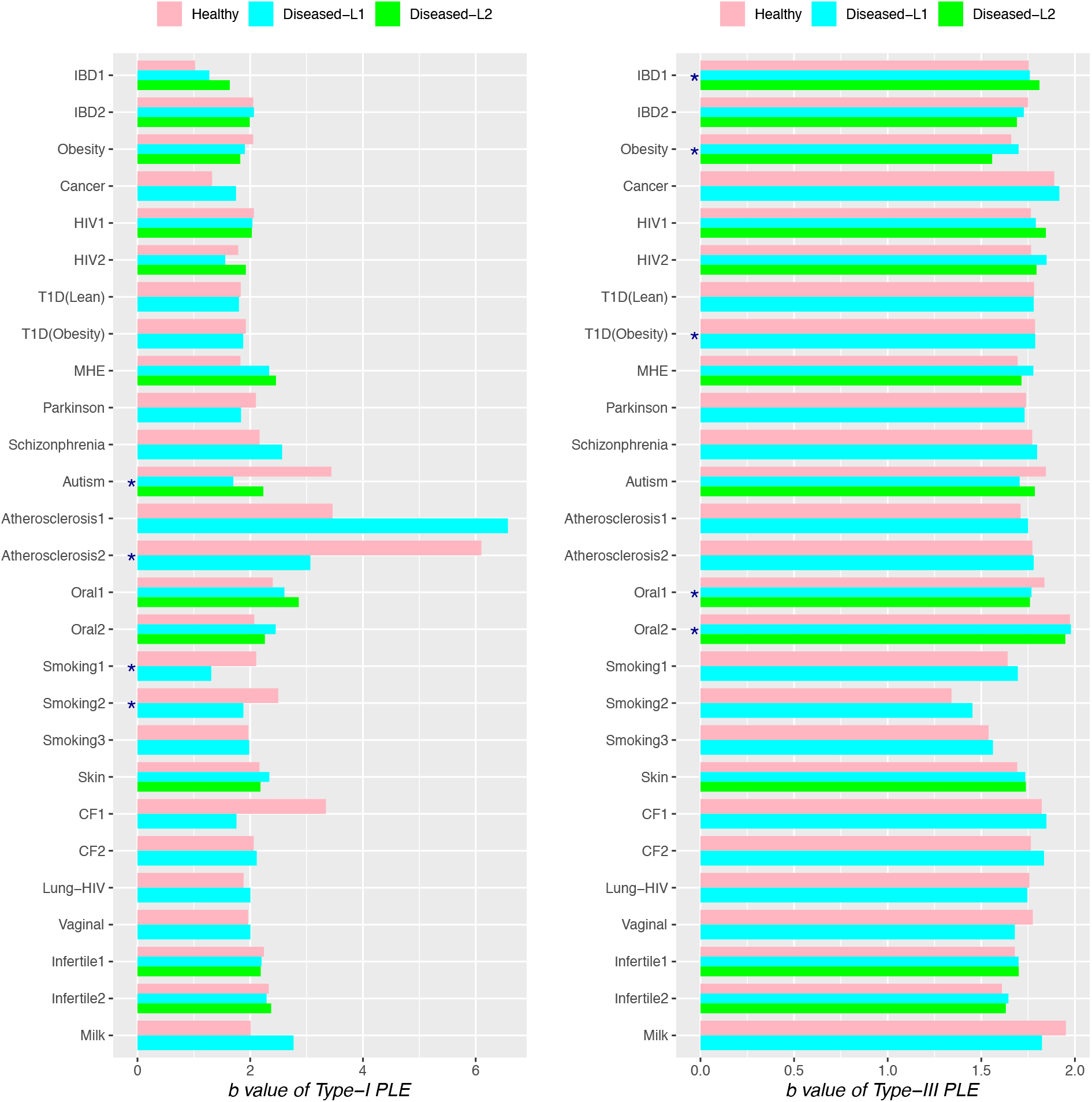
Bar graphs displaying the results from randomization tests for the difference in the heterogeneity scaling parameter (*b*) between the *H* (healthy) and *D* (diseased) treatments, the MAD (microbiome associated disease) cases with significant differences are marked with star (*).

From Table S2-S4, Figs 1–3, we summarize the following four findings:

i. All but one treatment in the 25 MAD studies were fitted to Type-I & III TPLE models successfully (p<0.05). The only exception (the *H* treatment of case #1) exhibited ap-value of 0.067, which is not too far from the significance level (*p*=0.05) for determining the model fitting.
ii. The *observed* average community spatial heterogeneity parameter (*b*) of Type-I TPLE model in *H*treatment is 2.304 and in *D* treatment is 2.205. The observed average *b* of Type-III TPLE model in *H* treatment is 1.743 and in *D* treatment is 1.751. These parameter values were computed from the *observed H* & *D* treatment data without performing permutations, and they all fall within the normal range of the power law and its extensions (Ma 2015, Taylor 2019).
iii. Through the randomization tests for the differences in the heterogeneity parameter (*b*) between the *H* & *D* treatments, it was found that: Type-I TPLE (for measuring the *community spatial heterogeneity*) parameter *b* is different in 10% of the MAD studies; Type-III TPLE parameter *b* (for measuring *mix-species population aggregation)* is different in approximately 12.5% of the MAD studies.
iv. Through the randomization tests for the differences in the TPLE parameter (*a*) between the *H* & *D* treatments, it was found that: Type-I TPLE parameter *a* is different in 12.5% of the MAD studies; Type-III TPLE parameter *a* is different in approximately 17.5% of the MAD studies. This and the previous findings suggest that MADs appear to have a more far-reaching influence at the mixed-species level than at the community level.

## Discussion

In the below, we further discuss the implications of the four findings, illustrated above, from the perspective of the heterogeneity-disease relationship (HDR). Our discussion is approached from comparison with our previous diversity-disease relationship (DDR) study (Ma *et al*. 2019).

First, the findings from the above HDR analysis echoed the conclusions from our previous DDR study (Ma *et al*. 2019). In the previous study, the diversity in the Hill numbers (Chao *et al*. 2014) exhibited significant differences between the *H* & *D* in only approximately 1/3 of the MAD cases, and here the percentage from HDR analysis ranged between 10%-12.5%, which is less than ½ of the percentage with difference from previous DDR analysis. This suggests that the MADs as disturbances on the human microbiomes can only exert limited influences on the community heterogeneity and/or diversity scaling (changes), and the influences on the heterogeneity appear to be less than on the diversity. In other words, the human microbiome ecosystem possesses significant stability (resilience).

However, in terms of the individual MAD case, the congruency between previous DDR analysis (Ma *et al*. 2019) and HDR analysis in this study may or may not be aligned with each other. In the case of TPLE-I, there are two case studies (#14 & #18), which show significant disease effects in HDR analysis and also exhibited significant diseases effects in previous DDR analysis (Ma *et al*. 2019). However, there are two case studies (#12 & #17) that show significant disease effects in HDR analysis but exhibited no significant disease effects in previous DDR analysis (Ma *et al*. 2019). In the case of TPLE-III, the congruency between previous DDR analysis and HDR analysis in this study is aligned fully with each other indeed. All four MAD studies (#2, #4, #15 & #16) that exhibit significant disease effects in the HDR also showed significant disease effects in previous DDR analysis (Ma *et al*. 2019).

Second, as to the reason why HDR analysis reveals a less significant percentage of differences (disease effects) than previous DDR analysis, the answer can be found by distinguishing the nature of the *diversity metrics* (Hill numbers) and the TPLE model (heterogeneity parameter b). What DDR analysis reflects are primarily the *ecological* characteristics. In contrast, what HDR analysis, particularly Type-I TPLE, reflects are primarily the *evolutionary* characteristics. That is, the parameter (*b*) of TPLE is an evolutionary characteristic, which is well documented in existing literature (Taylor 1961, 1980, 1981, 1984, 1986a, 1986b, 2019, Taylor & Taylor 1977, Taylor *et al*. 1983, 1988). Although, especially for microbes, the evolutionary and ecological characteristics are often interwoven with each other and a clear division is usually difficult, the evolutionary properties should be more robust (stable) in general. This explains the relatively lower disease effects on the heterogeneity than on the diversity because the heterogeneity as evolutionary property should be more robust (stable).

Third, as to the reason why TPLE heterogeneity parameter (*b*) showed a less significant percentage of differences (disease effects) than the other TPLE parameter (*a*) in detecting the MAD effects, the answer also lies in the fact that parameter (*a*) is not an evolutionary characteristic, instead, *a* is primarily influenced by sampling schemes (Taylor 1961, 1984, 1986a, 1986b, 2019). For this reason, parameter (*a*) is often attached less importance or even ignored in literature.

Finally, as to the reason why Type-III TPLE or the TPLE for mixed-species population aggregation showed a more significant percentage of disease effects than Type-I TPLE, the answer lies in the fact that Type-III TPLE is essentially still a population level model, although it uses community level data. This finding also mirrored the results from *shared species analysis* in our previous study (Ma *et al*. 2019). The shared species analysis in that study revealed that in approximately 50% of the MAD cases, the shared species between the *H* & *D* samples were reduced due to the disease effects. In other words, the disease effects are more significant at species scale than at community scale.

## Acknowledgements

This work was supported “Industry Technology Talent Grant”, and “A China-US International Collaborative Project” from Yunnan Science and Technology Bureau, of China.

## Author Contributions

ZS Ma designed the study, interpreted the results and wrote the paper. LW Li performed the computation and participated in the interpretation of the results.

## Data accessibility

All datasets analyzed in this study are available in public domain and the data sources for downloading are provided in the online supplementary Table S1.

## Compliance with ethical standards

N/A

## Conflict of interest

The authors declare that they have no conflict of interest.

## Online Supplementary Tables

**Table S1**. Twenty-six 16S-rRNA metagenomic datasets utilized to analyze the effects of microbiome-associated diseases (MADs) on the microbiome heterogeneity

**Table S2**. The parameters of Type-I power law extension (TPLE-I) for the community spatial heterogeneity for each *H* (healthy) or *D* (diseased) treatment of the 25 MAD studies

**Table S3**. The parameters of Type-III power law extension (TPLE-III) for mixed-species population spatial aggregation for each *H* (healthy) or *D* (diseased) treatment of the 25 MAD studies

**Table S4**. The *p*-values from the permutation tests (with 1000 times of permutation re-sampling) for the differences in the TPLE-I and TPLE-III parameter (b) between the *H* and *D* treatments

